# Barriers and incentives for conducting research amongst the Ophthalmologists in Sub-Sahara Africa

**DOI:** 10.1101/321679

**Authors:** Kazim A Dhalla, Micheal Guirguis

## Abstract

**Background:** Research is a critical component amongst the strategies to improve health outcomes of any country. The role of research assumes greater importance in Africa as it carries a larger share of global burden of diseases, blindness and low vision. “Vision 2020- the Right to Sight” is a WHO-IAPB collaborated initiative aiming to eliminate preventable blindness by the year 2020. High quality research in eye care is imperative for the initiative to succeed, however, there is a dearth of research in eye care in sub Saharan Africa in general and specifically in the Eastern, Central and Southern African (ECSA) region. Identifying the barriers that hamper research in this region is an important step towards elimination of preventable blindness.

**Methods:** A structured questionnaire using the SurveyMonkey program was sent to ophthalmologists in the ECSA region and South Africa through their respective regional professional bodies. Data was analyzed using the SPSS program version.

**Results:** Lack of funding, inadequate time and poor research knowledge were the main research barriers while ability to improve eye health care through research was the main incentive for conducting research.

**Conclusion:** The barriers mainly center on financial, human and administrative infrastructure and resources. In spite of the barriers, ophthalmologists in the study region are enthusiastic in research aiming to increase evidence based knowledge to improve eye health care in line with the goals of “Vision 2020- the Right to Sight” initiative.

## Introduction

Africa carries a large burden of global blindness and visual impairment. By the WHO estimates, 60% of the world’s blind live in Sub Saharan Africa, India and China. In 1999, WHO in partnership with International Agency for the Prevention of Blindness (IAPB) launched a global initiative called “Vision 2020- the Right to Sight” targeting to eliminate avoidable blindness, which is preventable in 80% of the cases, by the year 2020. (http://www.who.int/mediacentre/factsheets. Research would therefore be an integral part of this initiative if it were to achieve its goals. Though India and China are thriving in ophthalmic research (1), Africa is lagging behind to a large extent (2) with obvious paucity of scientific literature from Sub Saharan Africa in the major medical databases (3). Why is there low research productivity in Sub Saharan Africa? If barriers for conducting ophthalmic research exist, with the exception of West Africa, they are unknown for a large part of Sub Saharan Africa. This study explores the barriers and incentives for conducting research amongst the ophthalmologists in Sub Saharan Africa with specific focus on ophthalmologists in the Eastern, Central and Southern African countries (ECSA) and South Africa.

### Aim

To identify factors that act as barriers for conducting research and factors that encourage research activities amongst the Ophthalmologists in the ECSA region and South Africa.

### Methodology

Cross sectional survey of Ophthalmologists in the ECSA region (which is formed by the following countries; Tanzania, Kenya, Uganda, Rwanda, Burundi, Democratic Republic of Congo (DRC), Ethiopia, South Sudan, Zambia, Malawi, Botswana, Mozambique, Somalia and Lesotho) and South Africa (SA). West Africa was excluded from the study because a similar study was conducted in the region in 2011 (4)

#### Study design

Cross sectional survey study

#### Inclusion criteria

All ophthalmologists in the ECSA region and SA irrespective of ethnicity and whether in clinical practice, research, administration or retired.

#### Exclusion criteria

African ophthalmologists originally from the study region but currently residing out of the study region.

#### Sample size

The study region is estimated to have about 622 Ophthalmologists and distributed as shown in “Table 1”:

**Table 1:**
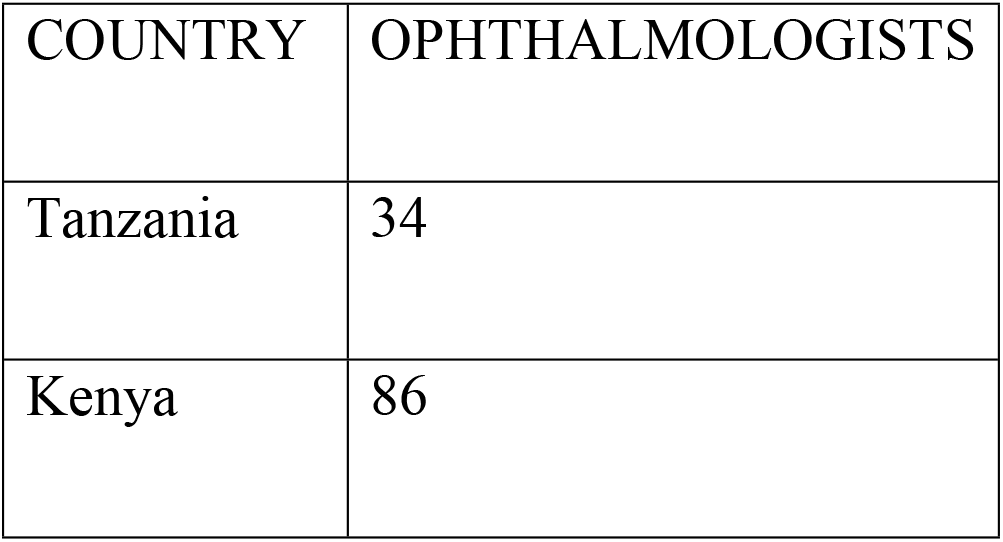
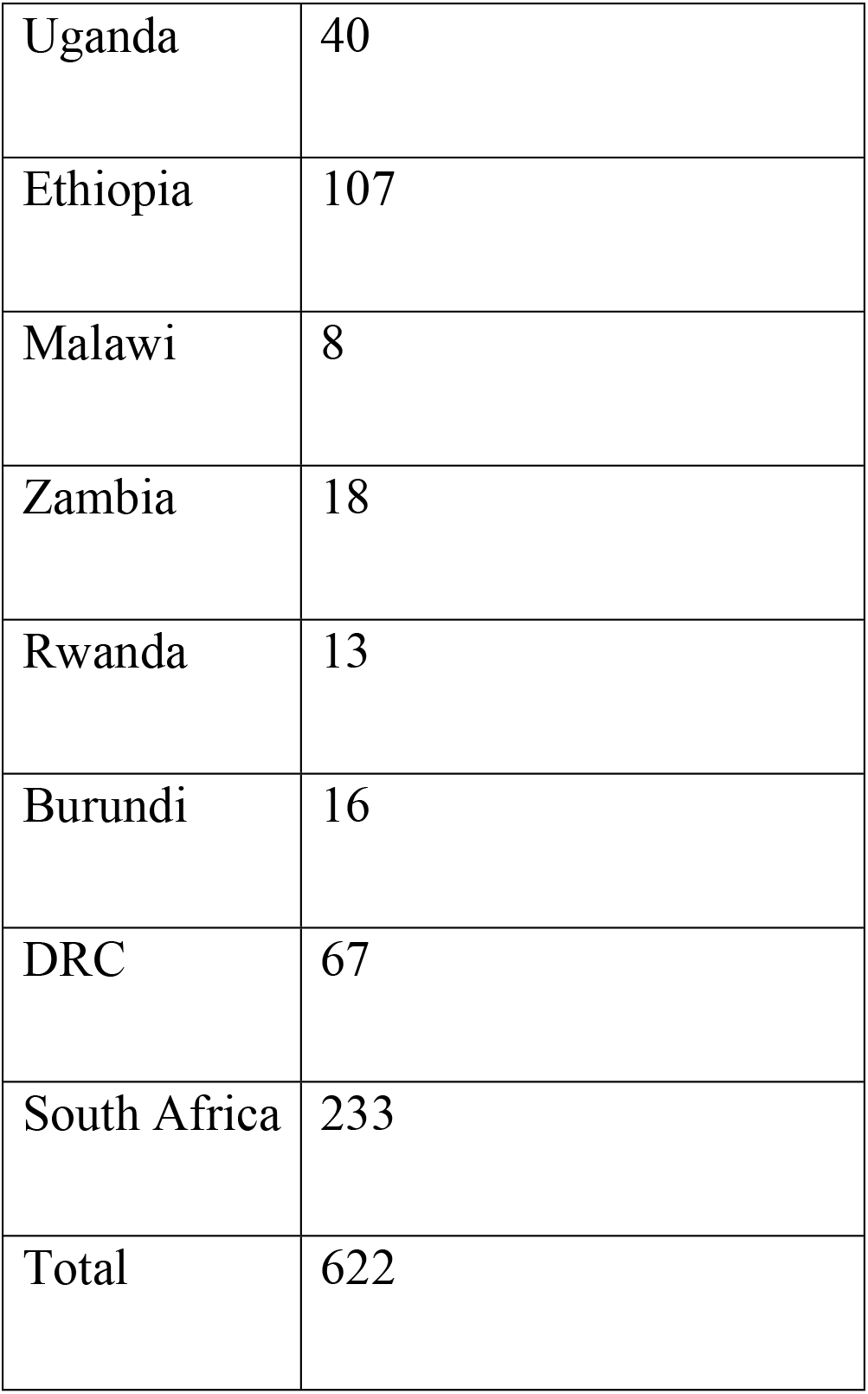
Distribution of Ophthalmologists in the study region.

(www.icoph.org. 2012 and (5)

Assuming 20% of the participants (124) are not reachable and response rate of 60%, the sample size was calculated to be 300 as follows:

622–124 = 498X 60% = 298.8 rounded to 300.

However, the target was to register all the ophthalmologists hence census sample was extracted.

#### Survey tool

A structured questionnaire was used to collect data from the participants using the online SurveyMonkey ^®^ program (www.surveymonkey.com). Some questions were adapted,with the author’s permission, from a similar study done in Nigeria, West Africa (Mahmoud, A. et al 2011). The full questionnaire is accessible from https://www.surveymonkey.com/r/Eye_Research

#### Pilot testing

A URL linked questionnaire was sent to 10 Ophthalmologists outside the study area and were requested to participate in the pilot study. All respondents found the questionnaire easy to fill and took less than ten minutes to complete. All but one thought that the questions were relevant to the objectives.

#### Ethical issues

The study is extracted from a dissertation for a Master of Science degree (M.Sc) in clinical research with University of Liverpool (online course). Ethical clearance for the study was granted by the University of Liverpool ethics committee after obtaining permission to conduct the research in the study area from the regional ophthalmological bodies; College of Ophthalmology of Eastern, Central and Southern Africa (COECSA) and Ophthalmological Society of Southern Africa (OSSA) respectively.

#### Participant recruitment procedure

The COECSA secretariat office sent out electronic mails with the URL link of the questionnaire to all the members. The study was also advertised on OSSA’s web based monthly newsletter circulated by electronic mail to all the members. Subsequently, 3 follow up reminders were sent to the members requesting them to participate in the study. Additionally, personal mails with two follow up reminders were sent to chairpersons of individual countries’ ophthalmological societies requesting them to encourage their members to participate. Participant recruitment started on 1^st^ April 2016 and access to the questionnaire was closed on the 20^th^ of May 2016. It was however not possible to know how many ophthalmologists actually got the information.

Consent was taken by asking participants to “tick” the consent box in the questionnaire if they agreed to participate. Furthermore, the action of filling the questionnaire itself was taken as surrogate for consent. Participant Information Sheet (PIS) was included in the online questionnaire which specified that participation was voluntary with the option of not answering personal questions like name, age and/or gender. Access to database was restricted to the researchers only thus ensuring complete confidentiality of the participants’ information.

#### Data entry process

All responses were stored in the SurveyMonkey^®^ program and directly downloaded to the statistical package, SPSS^®^ version 21 (IBM ^®^SPSS^®^Statistic) in the coded form.

### Data cleaning and analysis

Data cleaning was done by running a frequency distribution of all the variables. A few respondents preferred not to mention their names, age and/ or gender. Questionnaires without the demographic data were included in the analysis as this would not affect the overall results.

Research productivity, defined as the number of research papers published in the 10 year period from 1^st^ January 2005 to 31^st^ December 2014, was the dependent variable. Independent variables included age, gender, number of years in practice, type of institution i.e. government, non-government or private, post held i.e. Clinical, academic, both clinical and academic or purely administrative; time spent in private practice, additional post graduate training in research and pre-defined research barriers. Descriptive statistic was done using the frequency distribution and measure of central tendency appropriate for the data.

Inferential statistics was done using chi-square test for the 2 by 2 nominal variables and Somer’s delta (Somer’s d) test for the ordinal variables. Poisson regression analysis was done to determine statistical association between the dependent and independent variables. A p value of <0.05 was taken to be statistically significant.

## Results

### Survey response

There were a total of 114 respondents from the region. The response rate was therefore 38% assuming that survey questionnaire reached all the 300 potential participants. Country wise distribution of the respondents is given in “Table 2”.

**Table 2:**
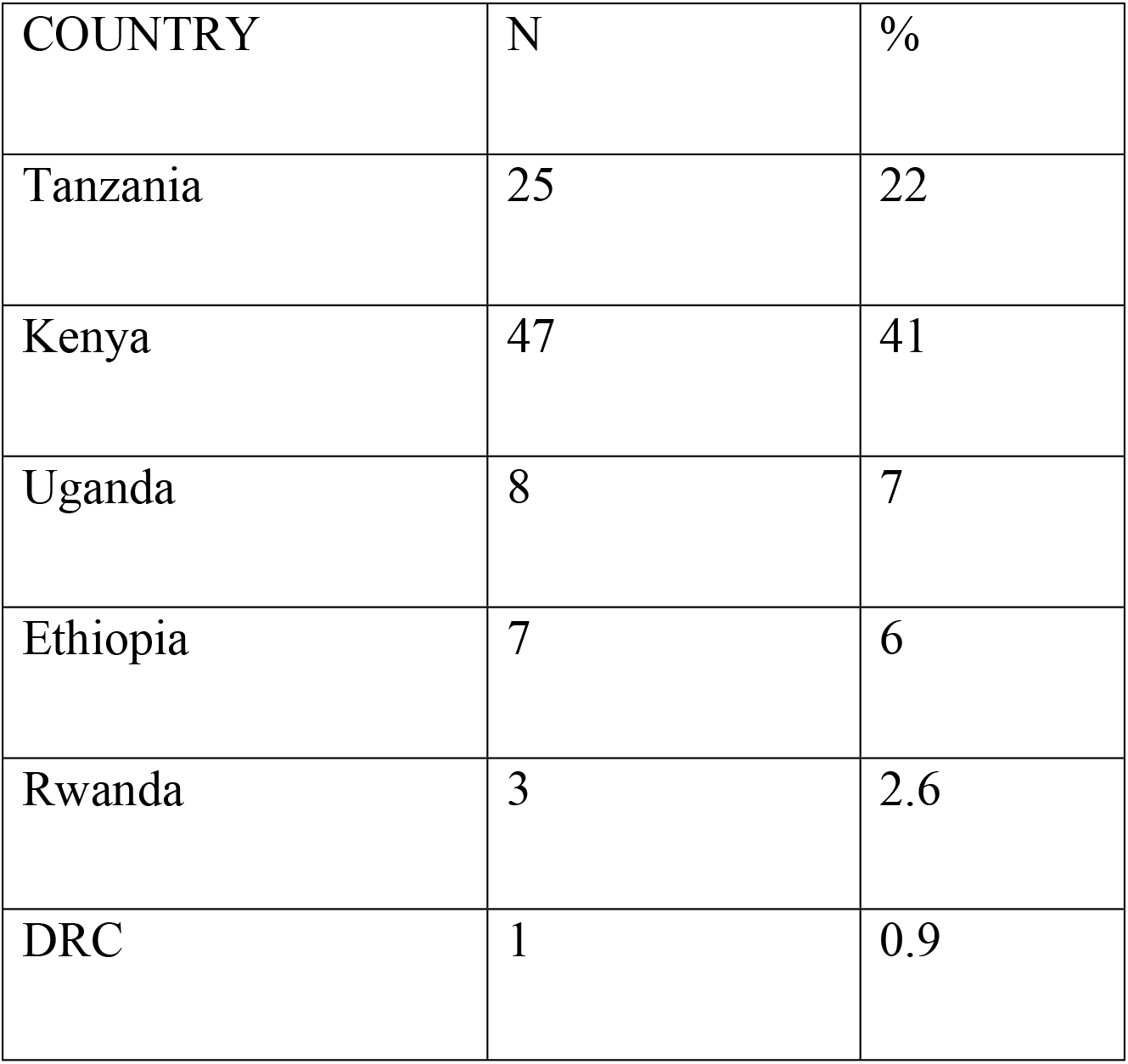
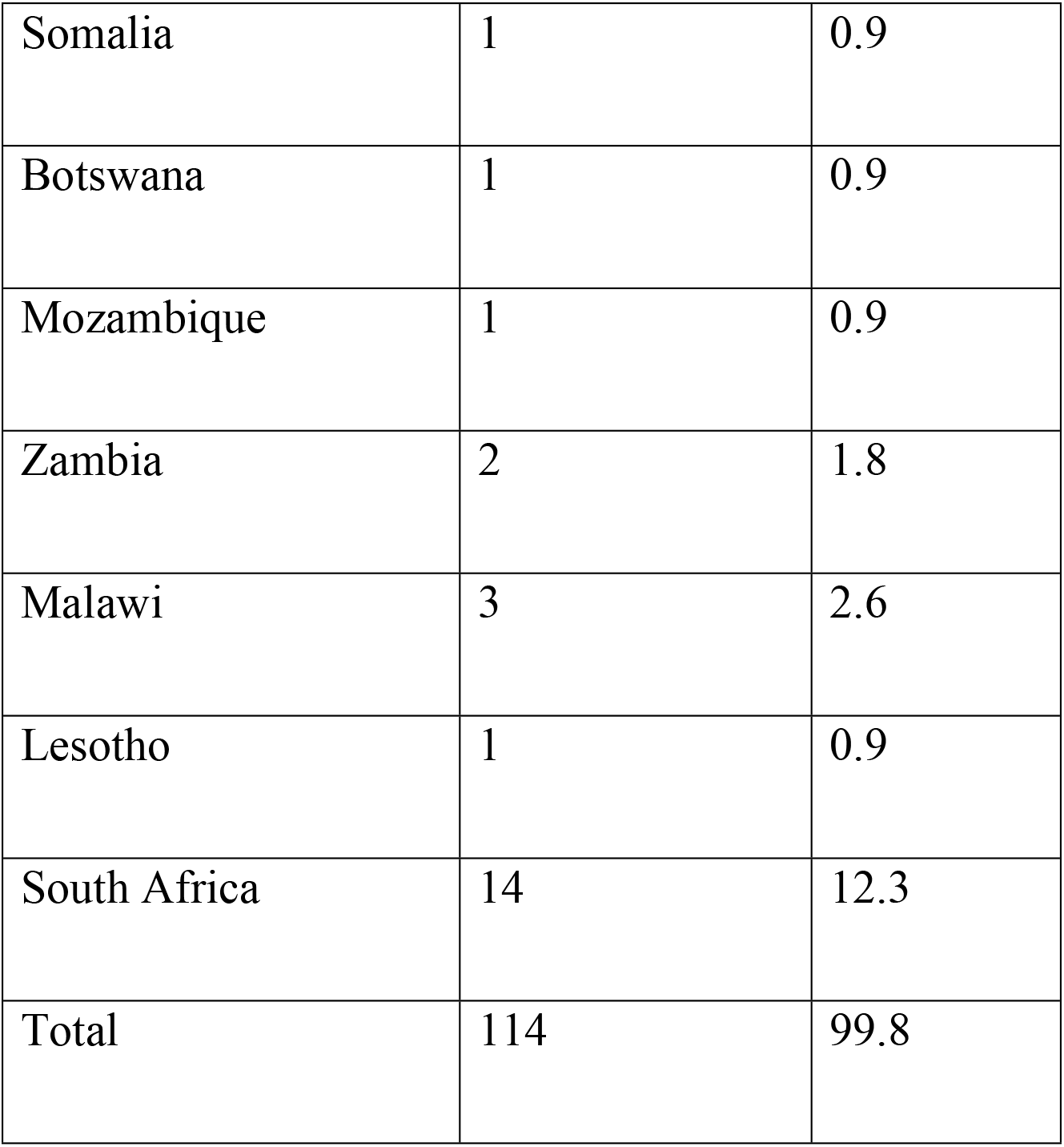
Distribution of respondents by study region.

Majority of the respondents, 87.7%, were from the ECSA region.

### Socio-Demographic description of the population

**Table 3:**
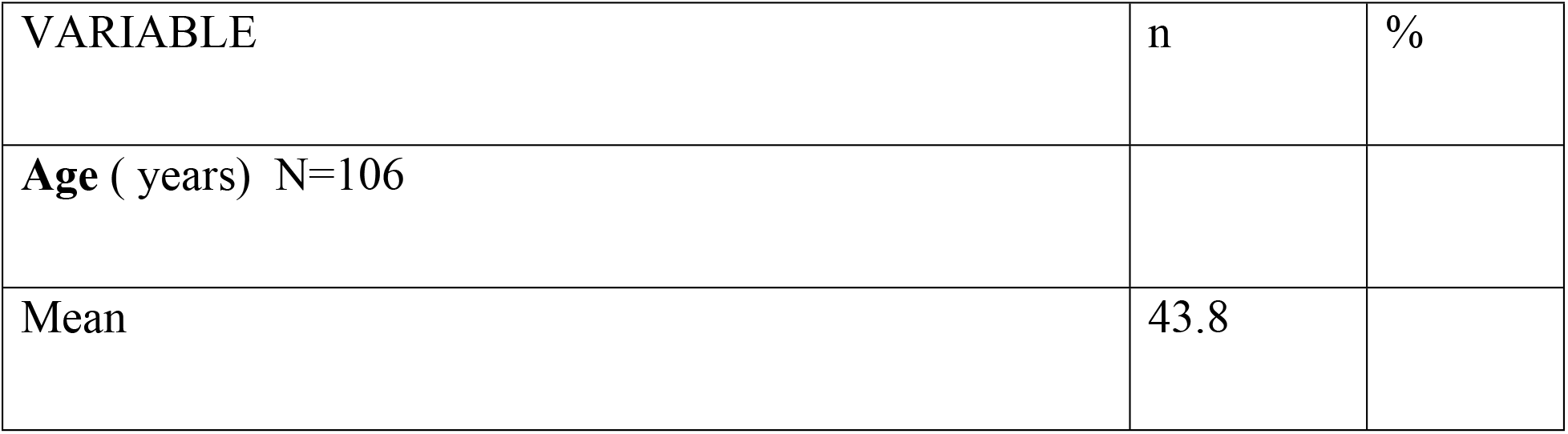
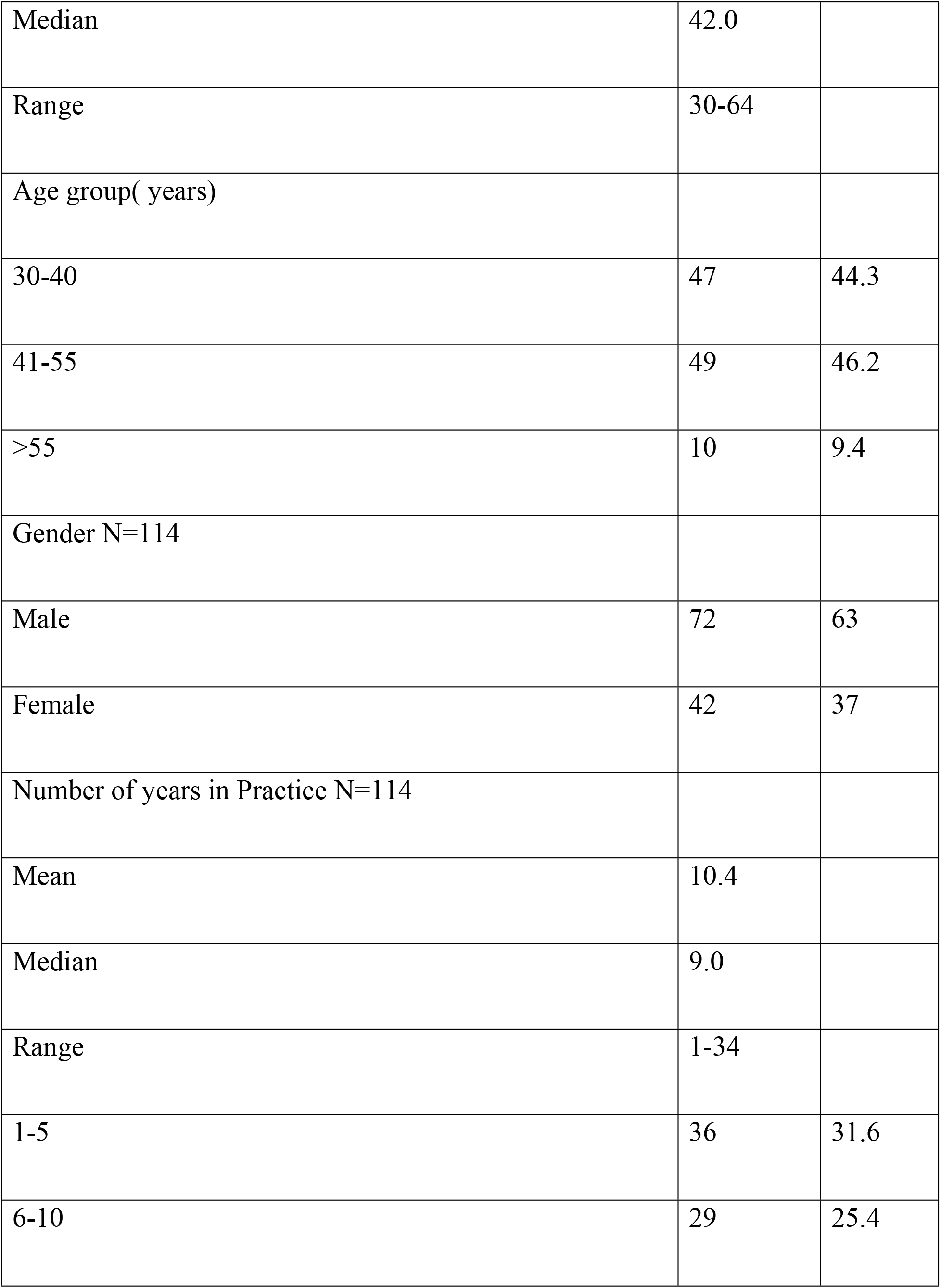
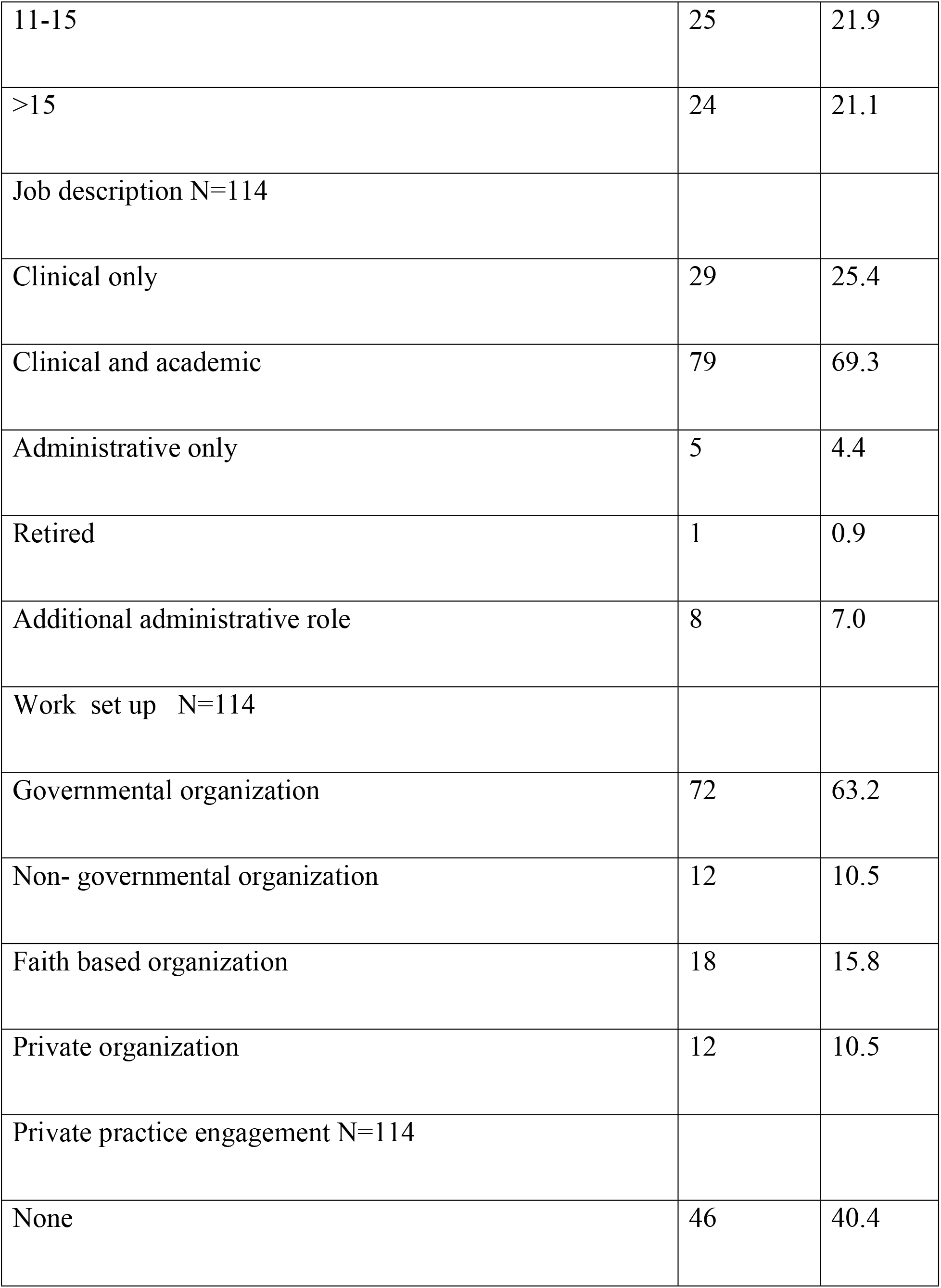
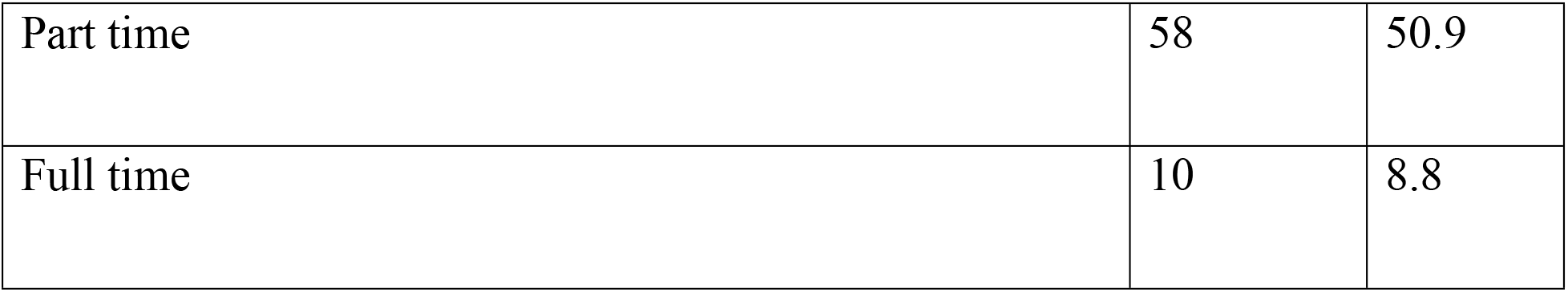
Socio-Demographic description.

A third of the respondents were in their early careers (1–5 years) in Ophthalmology and 19.3% were fresh graduates. Majority of the clinicians(69.3%) were also involved in academic practice either as university lecturers or teaching younger cadres including residents on attachment and cataract surgeons. 56 participants (49.12%) were practicing at only one place and that includes those who are fully engaged in their own private practice. The remaining 58 (50.87%) had full time placement as well as engaging in private practice.

### Academic and research profile

Respondents’ academic and research profiles and areas of research interest are given in “Table 4” and “Table 5”.

**Table 4:**
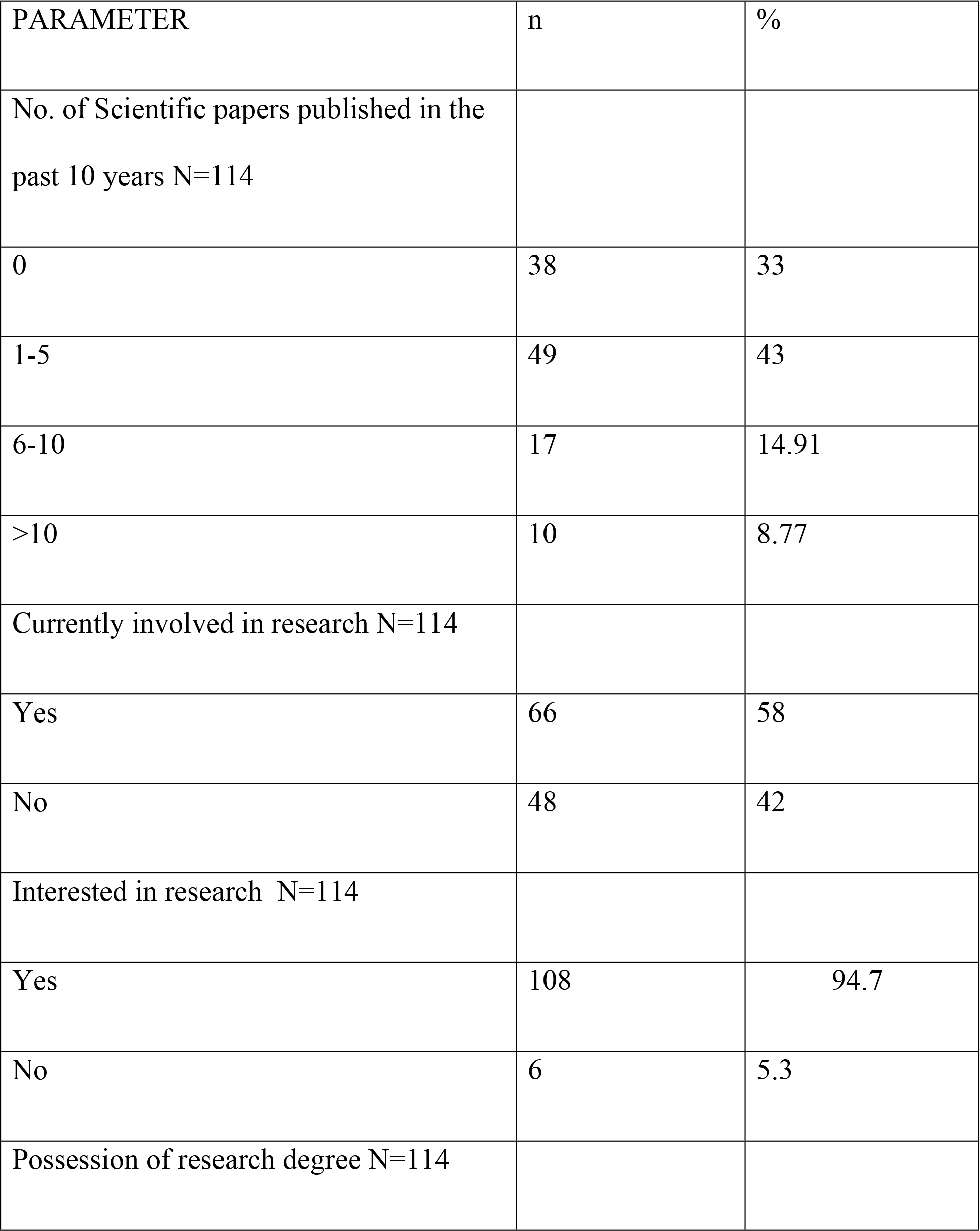
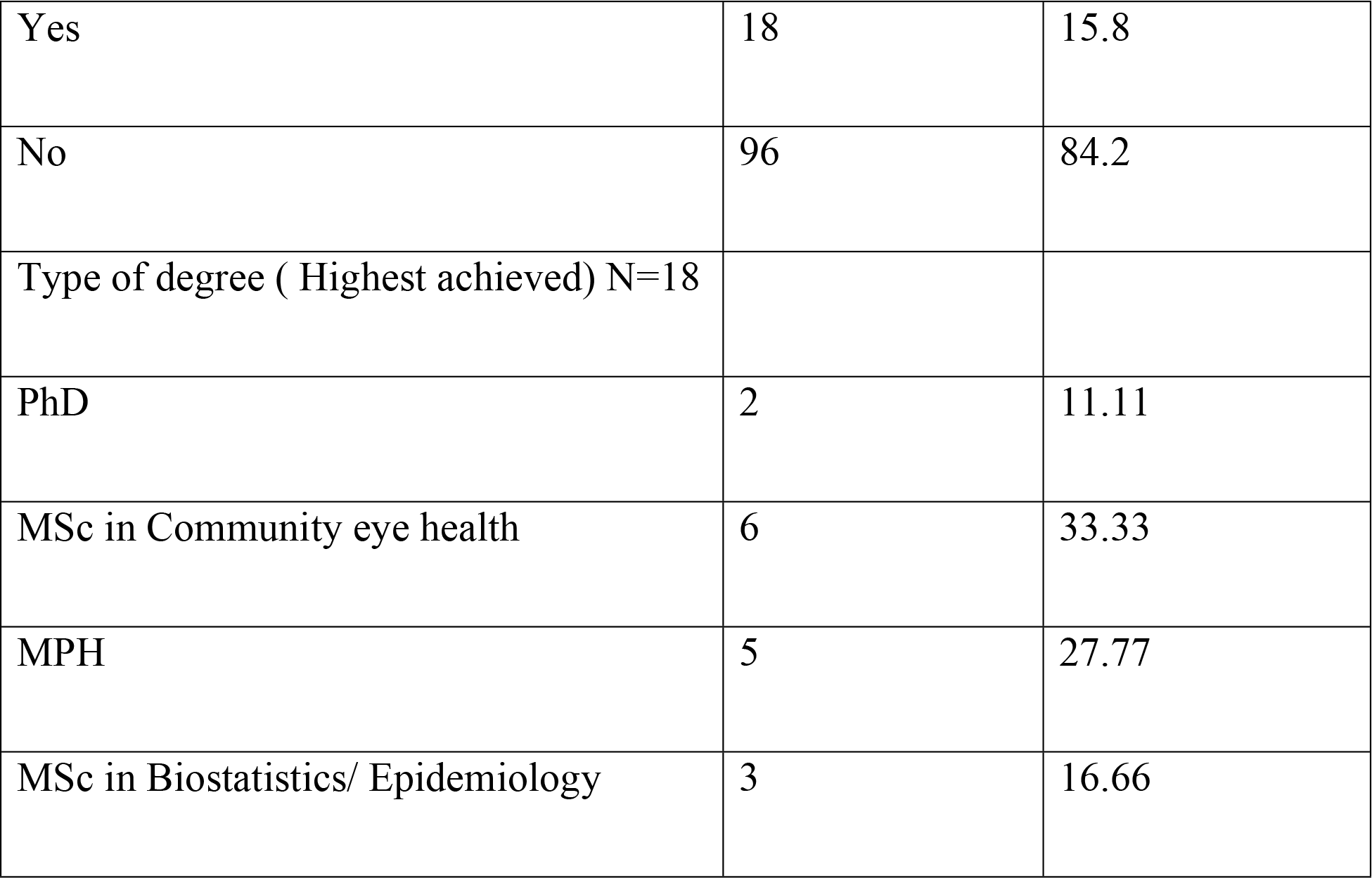
Academic and research profiles.

**Table 5:**
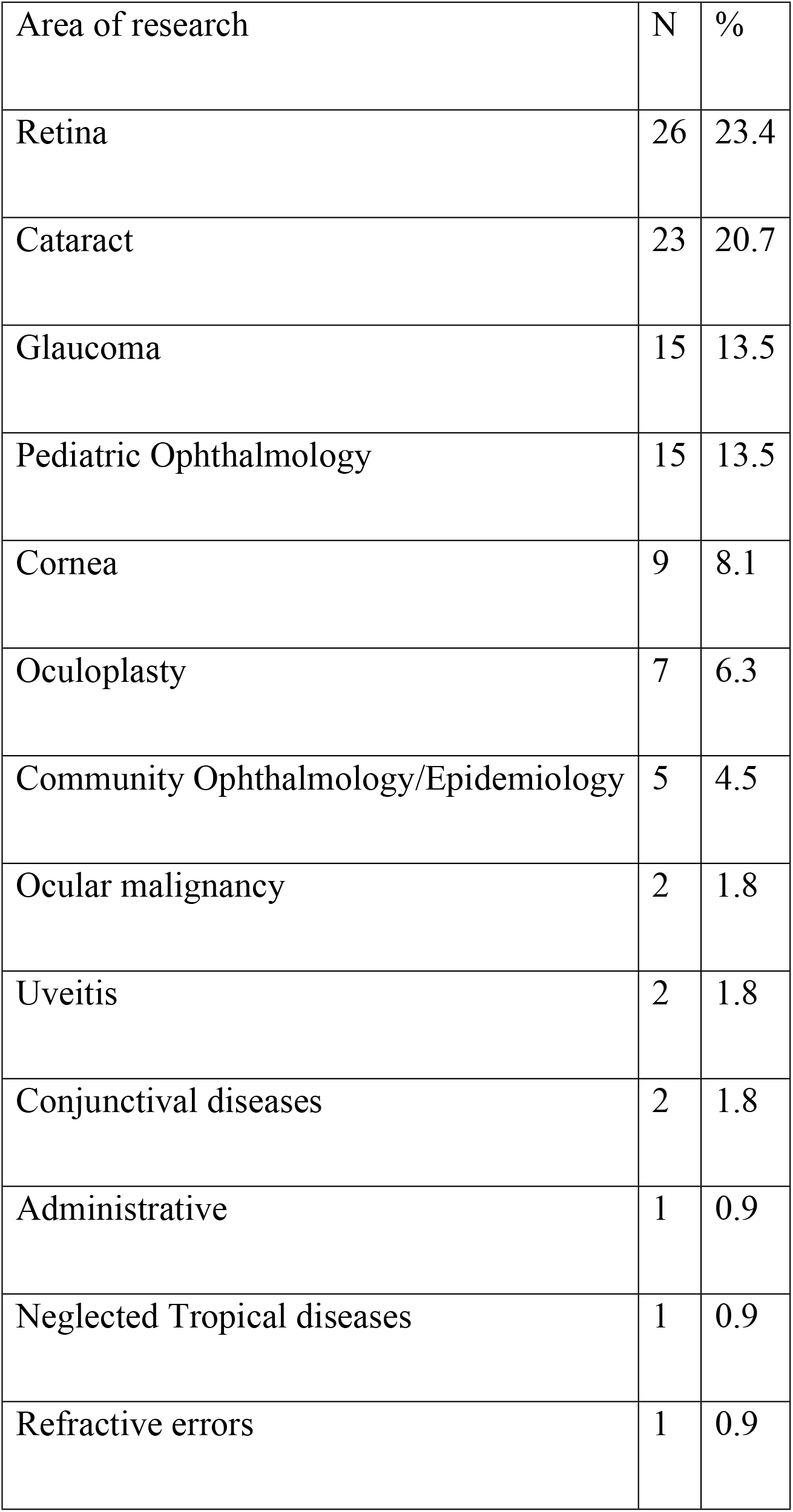
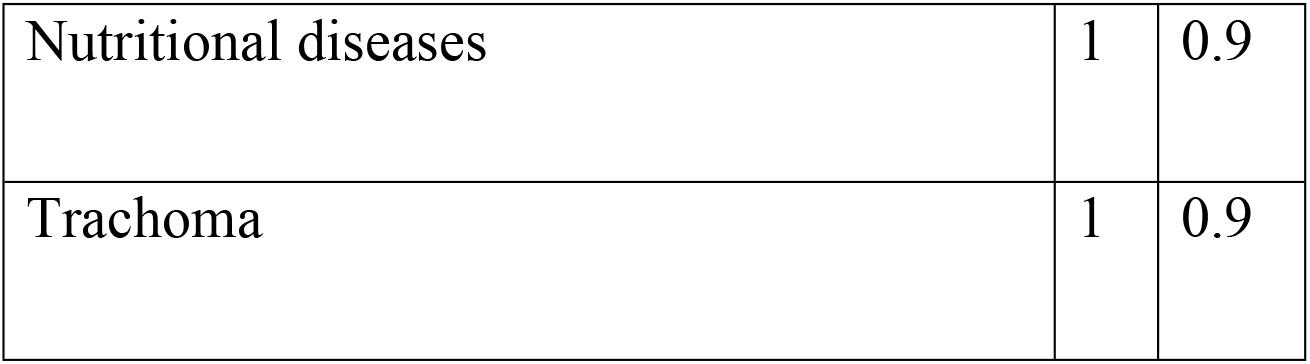
Broad areas of research interest.

A third of the respondents did not have any scientific publication in the past 10 years and 19 (16.7%) had only 1 publication. Number of papers published was statistically significantly related to the participant’s age (p = 0.000) and number of years in practice (p = 0.000), however, Spearman correlation coefficient was not very strong in both the cases; Age ρ = 0.437 and years in practice ρ = 0.5. There was no statistically significant association between the number of papers published and possession of a research degree (p = 0.077). 6 participants (5.2%) published at least 20 papers, all of whom had research degrees. One participant with PhD published 40 papers.

Retinal diseases had the highest research interest followed by cataract, glaucoma and pediatric ophthalmology. Though 14 participants had research degrees in community health and epidemiology, only 5 had interest in the field. It is interesting to note that only 1 respondent was interested in trachoma.

### Research barriers

The majority of respondents, 101/110 (91.8%), felt there were significant barriers for conducting ophthalmic research in Sub Saharan Africa. Figure 1 shows the frequency distribution of the barriers mentioned by the respondents.

**Figure 1:**
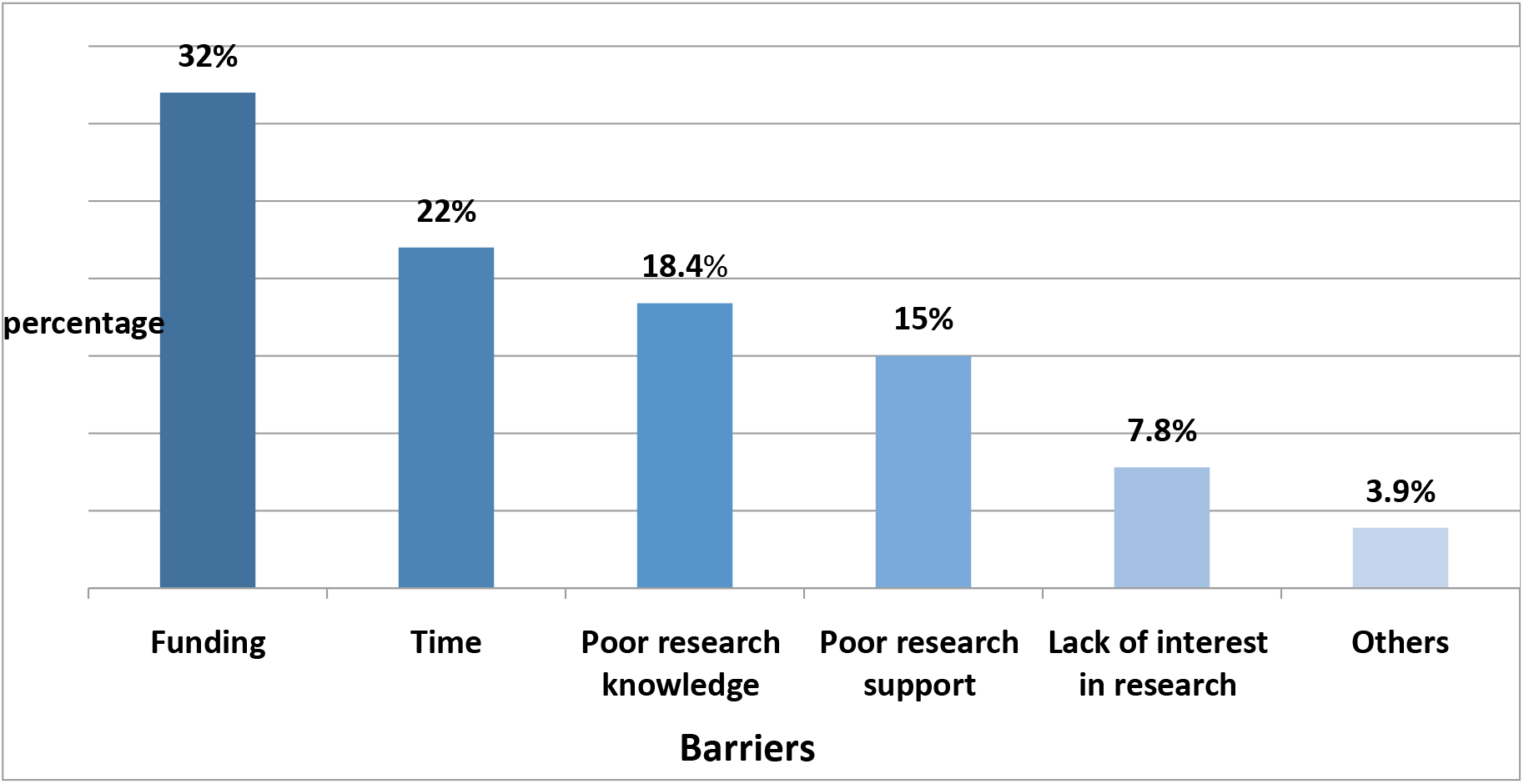
Research barriers.

### Funding

Majority of the respondents, 60 (53.6%), cited funding to be the major barrier for conducting research. Non-Governmental Organizations (NGO) stood out to be the most important source of research funding. Nearly a quarter of the respondents used personal funds for research while a third of the participants had never applied for research funding.

### Time

Half of the respondents had no time for research due to busy clinical commitments. A fifth had dedicated research time but majority thought that was not enough. There was no statistically significant relationship between availability of research time and whether one worked in a government or private set up, (p = 0.647).

### Knowledge

Respondents were asked to assess their knowledge in three broad areas; research process, common statistical software and word processing programs.

More than a half reported good knowledge of all stages of the research; however, statistical skills were poor in a large proportion of participants. Research knowledge was statistically significantly related to having an additional research degree (p = 0.001). ECSA participants reported significantly better statistical skills than the South African peers (p = 0.01). SPSS and EpiInfo were the two commonly known statistical programs, however, majority of the respondents had poor working knowledge of all the statistical packages.

### Research support

Research support was assessed on two areas, general research support given at the work place and access to electronic resources. Generally, respondents reported poor research support at their work places, however, research support is better in academic compared to non-academic institutions (p = 0.016) and access to ethical committees was better in government compared to non-government institutions (p = 0.000).

Though internet was readily available, e resources including HINARI was not accessible to the majority of the respondents. There were significant differences between ECCSA countries and South Africa in this area. E resources were more accessible to South African respondents (p = 0.045) while HINARI was more accessible to ECSA respondents (p = 0.003).

### Publication barrier

Majority of the respondents (75/108, 69.4%) felt it was difficult for the African researchers to publish in non-African journals. There was no difference between the ECSA and South African participants (p = 0.108)

### Factors associated with research output

A multivariate analysis was done using the Poisson regression model to determine the factors associated with research output. “Table 6” summarizes the results.

**Table 6:**
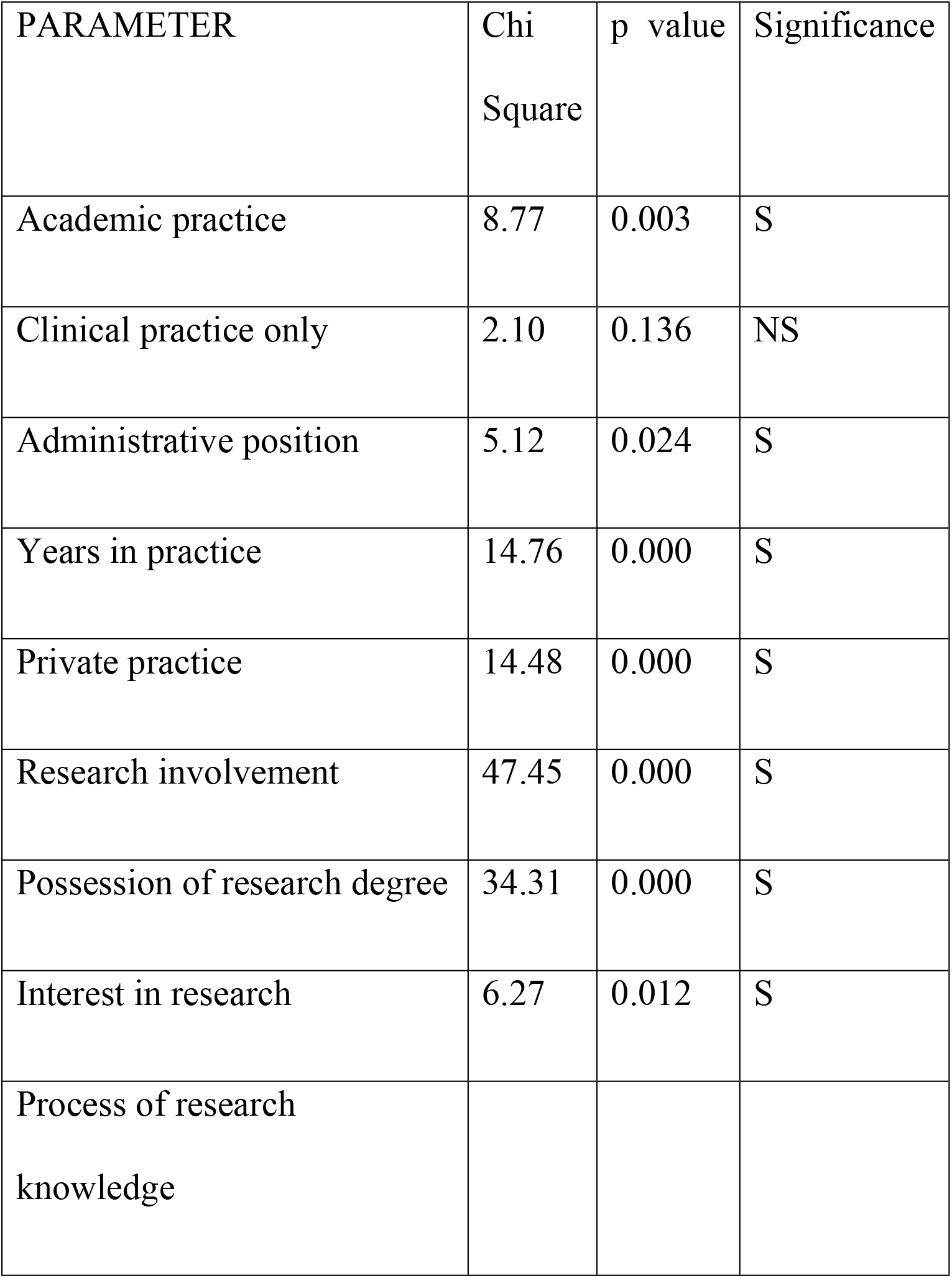
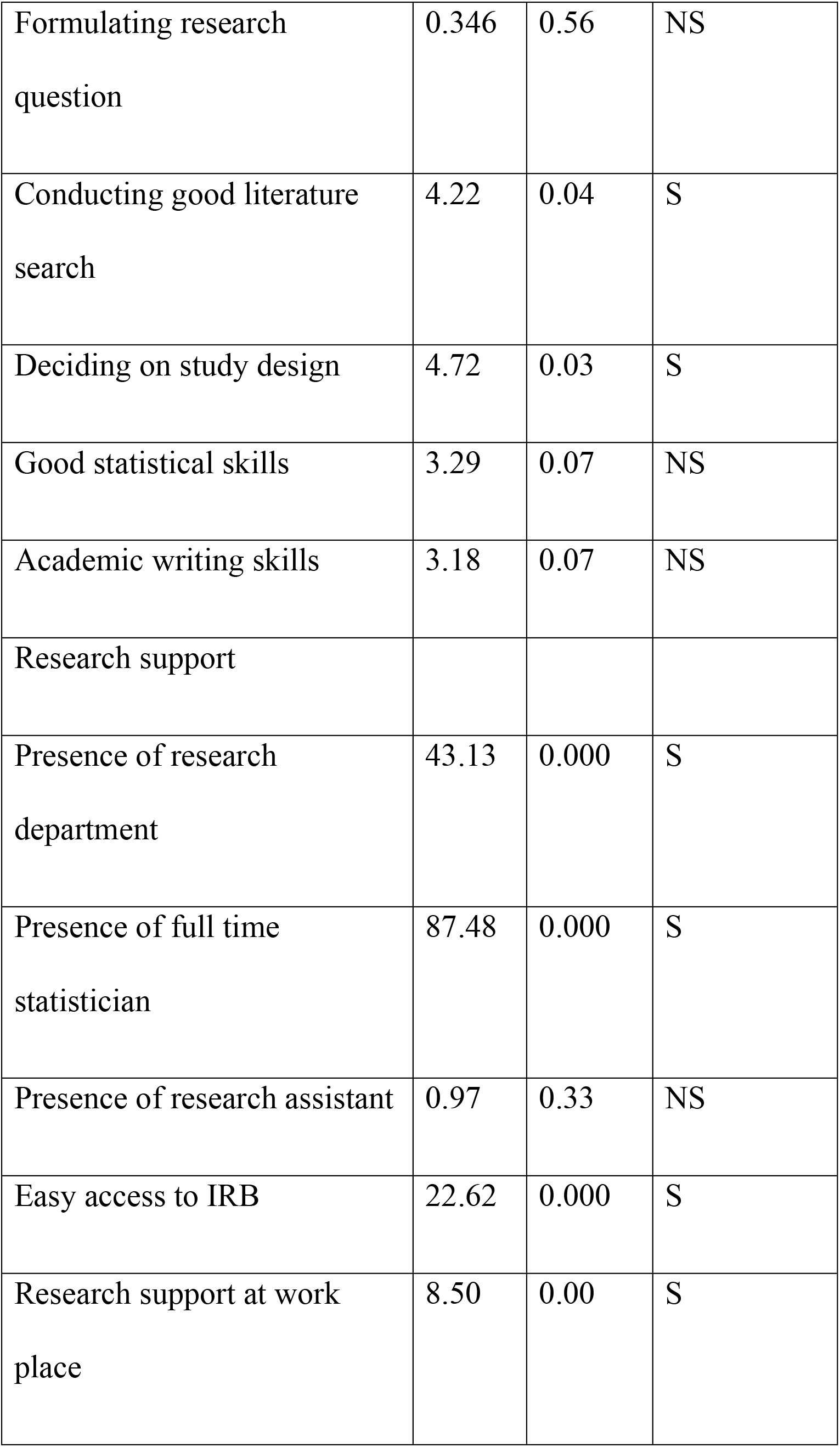
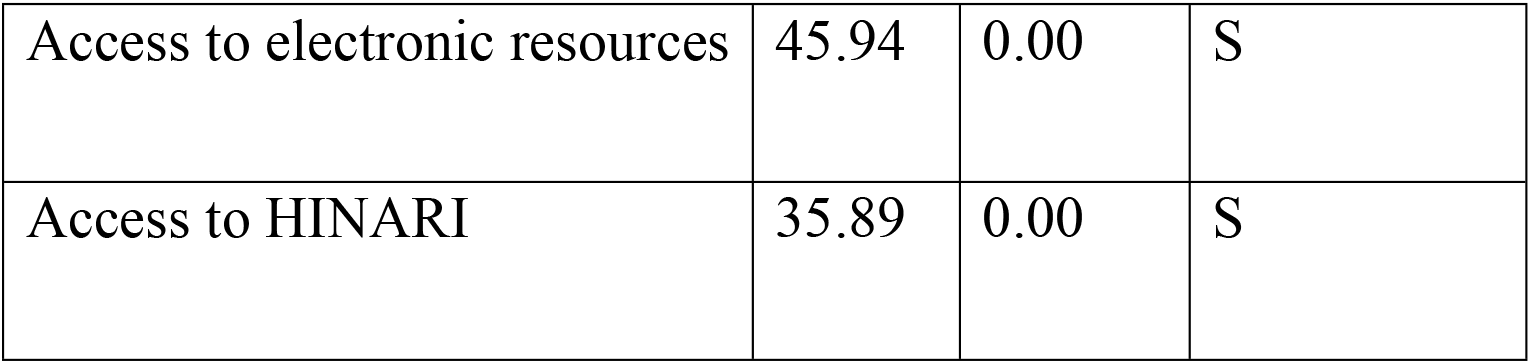
Factors associated with research output.

S = Factors which are statistically significantly associated with research output

NS = Factors which are not statistically significantly associated with research output.

Figure 2 gives a frequency distribution of the incentives that drive the participants to conduct research in the region.

**Figure 2:**
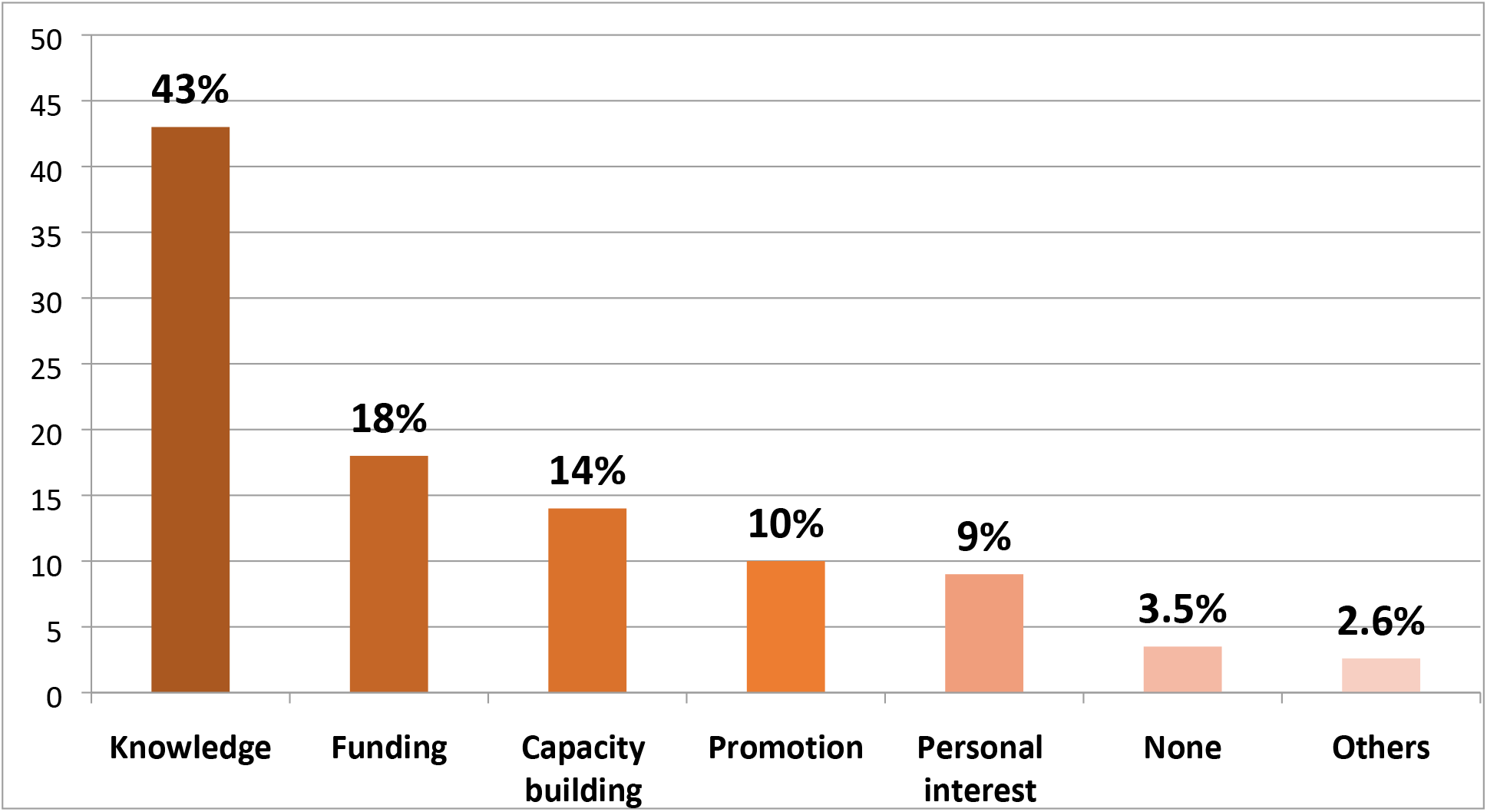
Incentives to conduct research.

The main incentive to conduct research was to expand the existing knowledge base in order to provide evidence based management of patients. A small proportion of respondents felt there were no incentives for conducting research.

## Discussion

A number of barriers and the associated factors for conducting ophthalmic research in the regions have been identified. Poor funding, inadequate time for research, poor research knowledge and departmental support were the prominent barriers. Research productivity was significantly associated with academic practice, possession of research degree, research knowledge, research support and access to electronic resources. Contributing to and expanding the existing knowledge base was the main incentive for conducting research amongst the ophthalmologists in the study region. There were no major differences in barriers and incentives between the ECSA and South African participants.

Most of the respondents were involved in research activities and a large majority was interested in research. The high interest in research amongst the ECSA ophthalmologist mirrors that of East African Orthopedic surgeons (6) but strikingly different to the poor research interest amongst the Nigerian Ophthalmologists (4). The authors of the Nigerian study felt that poor research interest amongst the Nigerian ophthalmologists is due to funding constraints and inadequate knowledge in research process which feature among the major barriers for conducting research in Nigeria.

### Research barriers

#### Funding

Research funding was one of the main barriers for conducting research. This is consistent with the findings from Nigeria amongst the ophthalmologists (7) and medical specialists (8) and East African orthopedic surgeons (6). Personal funds was the main source of funding for Nigerian ophthalmologists and medical specialist. Research is an expensive venture and if researchers have to rely on personal funds for research then this is a great disincentive for conducting good quality, high impact research which also requires funds for publication. It appears that African researchers have not yet explored funding partnerships with the pharmaceutical industry or other corporate sponsors. Standard chartered bank for example is leading in funding eye care services and training in their novel Seeing is Believing (SiB) project in collaboration with a number of NGOs like the Fred Hollows foundation, ORBIS and IAPB (9). African researchers rate government funding last on the list in spite of WHO recommending and governments endorsing the 2% health budget dedication to research.

#### Time

Time constraint featured prominently in our study and appears to be a common barrier across different specialties and regions (Sabzwari et al., 2009, Lloyd et al., 2004, Brocato and Mavis, 2005). Though it is not practical to separate clinical practice from research work in Africa, a certain number of hours per week could be allocated to clinicians for research work. It was expected that ophthalmologists working in government set ups would have dedicated and perhaps more time for research compared to those engaged in private practice. This did not show up in our study.

#### Knowledge

Knowledge of research process was a significant barrier in our study. Statistical skills appear to challenge a large section of ophthalmologists. Hence, the presence of a statistician in the department was statistically significantly associated with increased research productivity. Decision on selecting appropriate study design and paper writing skills were also a problem though to a lesser extent. Ophthalmologists with additional post graduate training in research had good knowledge in all the components of research process and higher research productivity (p = 0.000). This component does not feature very well in Nigerian studies, however, it appears to be a major barrier amongst the orthopedic surgeons in East Africa as well as Asian doctors. This may be an indicator that there is inadequate training of research process both at the undergraduate and postgraduate level. Research involvement in medical school appears to have a stronger influence in research productivity (10).

#### Access to academic literature

Majority of the respondents had good internet facilities however most of the respondents did not have easy access to electronic academic literature and just about half had access to the Health InterNetwork Access to Research (HINARI) program. HINARI was initiated by WHO sponsored private –public partnership in 2002 and offers free access to a large collection of prestigious journals to health institutions in developing countries(11). In spite of this, 48% of the respondents in our study did not have access to it. This is in contrast to the Nigerian study where by electronic literature was the main source of scientific information to the ophthalmologists and HINARI is widely accessible to Nigerian researchers (12). A study on access to electronic scientific knowledge in selected East and West African countries found that more than a third of postgraduate doctors relied on textbooks for information and though internet was generally available, accessibility varied in private and national institutes. Generally awareness to free online resources including HINARI was low in West African compared to East African institutions. HINARI requires Institutional password and not accessible to individual researchers (13).

#### Research incentives

The biggest incentive for conducting research amongst the ophthalmologists in our study was to increase evidence based knowledge in the region. Another important incentive in our study was the ready availability of research funding which contrasts with the idea of actually looking for funding from donors. Research capacity building and academic promotions also featured as important incentives. Financial gain, fame and international travel to attend and present research findings did not feature at all in our study. It appears that ophthalmologists in this study are fully aware of the fact that research is not a venture for financial growth. Perhaps, there is also an element of altruism as well. Enhancement of knowledge was also the greatest incentive for conducting research amongst the Nigerian ophthalmologist and medical specialists, East African orthopedic surgeons and Asian doctors. However, financial gains and fame featured quite prominently in the Nigerian and Asian studies. Capacity building featured as the third most frequent incentive cited which parallels poor research knowledge as a barrier to research productivity. Building research capacity by training the local experts in research process appears to be a single most important factor that will address both barriers and incentives for research productivity (14).

## Conclusion

A number of barriers have been identified in this study which appear to hamper research productivity in Sub-Sahara Africa. Dedicated research time, research funding and lack of appropriate skills are the main barriers which if addressed will increase research output in the region.

## Acknowledgement

We wish to thank sincerely all the Ophthalmologists who took time from their busy schedules to participate in our study. Special thanks to the following colleagues who facilitated the study in one way or the other; Dr. Ebrahim Matende, Dr. Cyprian Ntomoka, Dr. Muchai Gachago, Prof. Trevor Carmichael, Dr. Andrew Boliter, Dr. Emeritus Chibuga and Josiah Onyango.

## References

1. Courtright P, Faal HB. How can we strengthen ophthalmic research in Africa? Can J Ophthalmol. 2006;41(4):424–5.

2. Budenz DL, Bandi JR, Barton K, Nolan W, Herndon L, Whiteside-de Vos J, et al. Blindness and Visual Impairment in an Urban West African Population: The Tema Eye Survey. Ophthalmology (Elsevier). 2012;119(9):1744.

3. Courtright P, Seneadza A, Mathenge W, Eliah E, Lewallen S. Primary eye care in sub-Saharan African: do we have the evidence needed to scale up training and service delivery? Annals of tropical medicine and parasitology. 2013.

4. Mahmoud AO, Ayanniyi AA, Lawal A, Omolase CO, Ologunsua Y, Samaila E. Ophthalmic research priorities and practices in Nigeria: An assessment of the views of Nigerian ophthalmologists. Middle East African journal of ophthalmology. 2011;18(2):164.

5. Palmer JJ, Chinanayi F, Gilbert A, Pillay D, Fox S, Jaggernath J, et al. Mapping human resources for eye health in 21 countries of sub-Saharan Africa: current progress towards VISION 2020. Human Resources for Health. 2014;12(1):1–16.

6. Elliott IS, Sonshine DB, Akhavan S, Slade Shantz A, Caldwell A, Slade Shantz J, et al. What factors influence the production of orthopaedic research in East Africa? A qualitative analysis of interviews. Clin Orthop Relat Res. 2015;473(6):2120–30.

7. Mahmoud AO, Ayanniyi AA, Lawal A, Omolase CO, Ologunsua Y, Samaila E. Survey of the attitudes of nigerian ophthalmologists to and resources for ophthalmic research. Middle East Afr J Ophthalmol. 2012;19(1):123–8.

8. Mahmoud AO, Ayanniyi AA, Lawal A, Omolase CO, Ologunsua Y, Samaila E. Survey of medical specialists on their attitudes to and resources for health research in Nigeria. Ann Afr Med. 2011;10(2):144–9.

9. Bank SC. Seeing is Believing. http://wwwstandardcharteredcom/. 2015.

10. Lloyd T, Phillips BR, Aber RC, Lloyd T, Phillips BR, Aber RC, et al. Factors that influence doctors’ participation in clinical research. Medical Education. 2004;38(8):848.

11. Aronson B, Aronson B, Aronson B, Aronson B, Aronson B. WHO’s Health InterNetwork Access to Research Initiative (HINARI). Health Information & Libraries Journal. 2002;19(3):164.

12. Anyaoku EN, Anunobi CV, Anyaoku EN, Anunobi CV, Anyaoku EN, Anunobi CV, et al. Measuring HINARI use in Nigeria through a citation analysis of Nigerian Journal of Clinical Practice. Health Information & Libraries Journal. 2014;31(2):148.

13. Helen Smith HB, Oscar Mukasa, Paul Snell, Sylvester AdehNsoh, Selemani Mbuyita, Masanja Honorati, Bright Orji, Paul Garner. Access to electronic health knowledge in five countries in Africa: a descriptive study. BMC Health Services Research. 2007;7(72):7.

14. Sawyerr A. African Universities and the Challenge of Research Capacity Development. Journal of Higher Education in Africa 2004;2(1):213–42.

